# TREM2 and APOE do not modulate phagocytic clearance of dying cells in the live mammalian brain

**DOI:** 10.1101/2023.03.17.533222

**Authors:** Eyiyemisi C. Damisah, Anupama Rai, Robert A. Hill, Lei Tong, Jaime Grutzendler

**Author notes:** contributed equally to this work (Eyiyemisi C. Damisah, Anupama Rai, Robert A. Hill).

## Abstract

TREM2 and APOE are two major risk factors for Alzheimer’s disease (AD) that have been proposed to play crucial roles in microglia pathophysiology by affecting their ability to phagocytose cellular debris or aggregated proteins. In this study, we investigated for the first time the impact of TREM2 and APOE on the removal of dying neurons in the live brain by implementing a targeted photochemical method for programmed cell death induction combined with high-resolution two-photon imaging. Our findings showed that the deletion of either TREM2 or APOE did not affect the dynamics of microglia engagement with dying neurons or their efficiency in phagocytosing corpses. Interestingly, while microglia that encapsulate amyloid deposits were capable of phagocytosing dying cells without disengaging from plaques or moving their cell bodies; in the absence of TREM2, microglia cell bodies were observed to readily migrate towards dying cells, further disengaging from plaques. Our data suggest that TREM2 and APOE variants are unlikely to increase risk of AD through impaired corpse phagocytosis.

**Summary:** High-resolution two-photon imaging of programmed cell death in the live mouse brain reveals that neither TREM2 nor APOE modulate microglia phagocytosis of neuronal corpses. However, TREM2 regulates microglia migratory behavior towards dying cells in the vicinity of amyloid plaques.

## INTRODUCTION

Cell death is a common occurrence in neurodegenerative diseases, including Alzheimer’s disease (AD)(Moujalled et al., 2021; Vila and Przedborski, 2003). Although the precise mechanisms of cell death in neurodegeneration are not fully understood, various forms of programmed cell death, such as necroptosis, apoptosis, and ferroptosis, have been implicated (Moujalled et al., 2021; Vila and Przedborski, 2003). Programmed cell death triggers a rapid and coordinated response from adjacent microglia and astrocytes aimed at efficient removal of cellular debris (Damisah et al., 2020). Mutations in phagocytosis-related genes, such as MERTK, as well as aging, can lead to inefficient debris clearance. Mutations in microglia-specific genes, such as CSF1r (Rademakers et al., 2011) or TREM2 (Kleinberger et al., 2014), are associated with neurodegenerative processes in the brain and retina (Gal et al., 2000).

In the brain, TREM2 is a microglia-specific receptor and a key genetic risk factor for AD. Trem2 has been implicated in microglia polarization towards amyloid plaques (Wang et al., 2016; Yuan et al., 2016), metabolic fitness (Ulland et al., 2017), and lipid sensing (Wang et al., 2015). It has also been linked to the phagocytosis of amyloid deposits (Kleinberger et al., 2014), myelin debris (Cignarella et al., 2020), and dying neurons (Hsieh et al., 2009). Recent studies have suggested that APOE, the main risk factor for AD, is a ligand for Trem2 (Atagi et al., 2015), and APOE-related signaling may mediate the switch from homeostatic to disease associated microglia (DAM) phenotype(Krasemann et al., 2017), potentially affecting the efficiency of debris phagocytosis. Thus, the Trem2-ApoE signaling axis may modulate diverse microglia responses in various neuropathological processes. However, direct evidence of the role of Trem2 or Apoe in phagocytosis and specifically in modulating the removal of dying cells in the live brain remains very limited.

To address this knowledge gap, we implemented novel methodologies to induce programmed cell death in the live brain using a targeted two-photon photochemical method (2Phatal) (Hill et al., 2017). This approach allowed us to obtain high-resolution time-lapse intravital images of the cell death and corpse removal processes for subsequent quantitative analysis. Using this methodology, we assessed the precise roles of TREM2 and APOE in the normal and AD brain to determine the extent to which they play modulating roles in the process of programmed cell death and corpse removal. Our data demonstrate that despite the many potential roles of TREM2 and APOE in microglia function and metabolism, deletion of these genes does not affect the dynamics of programmed cell death or the ability of microglia to engage with and remove dying cells in a timely and effective manner.

## RESULTS

### Strategy for investigating molecular modulators of microglial clearance of dying cells in the live mouse brain

In our previous work, we introduced a targeted photochemical method for inducing programmed cell death in individual cells in the live mouse brain (2Phatal) (Hill et al., 2017) (Fig 1A). This approach employs high spatiotemporal resolution two-photon imaging to examine the sequence of events and multicellular interactions that occur during cell death and corpse removal (Fig 1B). Our studies demonstrated that this technique can detect defective phagocytic phenotypes with high sensitivity, such as those associated with aging or deletion of phosphatidylserine receptors like MerTK (Damisah et al., 2020) (Fig 1C-E). In the present study, we used this method to investigate the influence of selected Alzheimer’s disease risk genes on microglial behavior during corpse clearance *in vivo*. Specifically, we applied this approach in AD-like mice and mice with Trem2 or Apoe deletion to quantitatively evaluate the effects of these genes on microglial engagement with dying neurons and their impact on corpse clearance efficiency.

**Figure 1:**
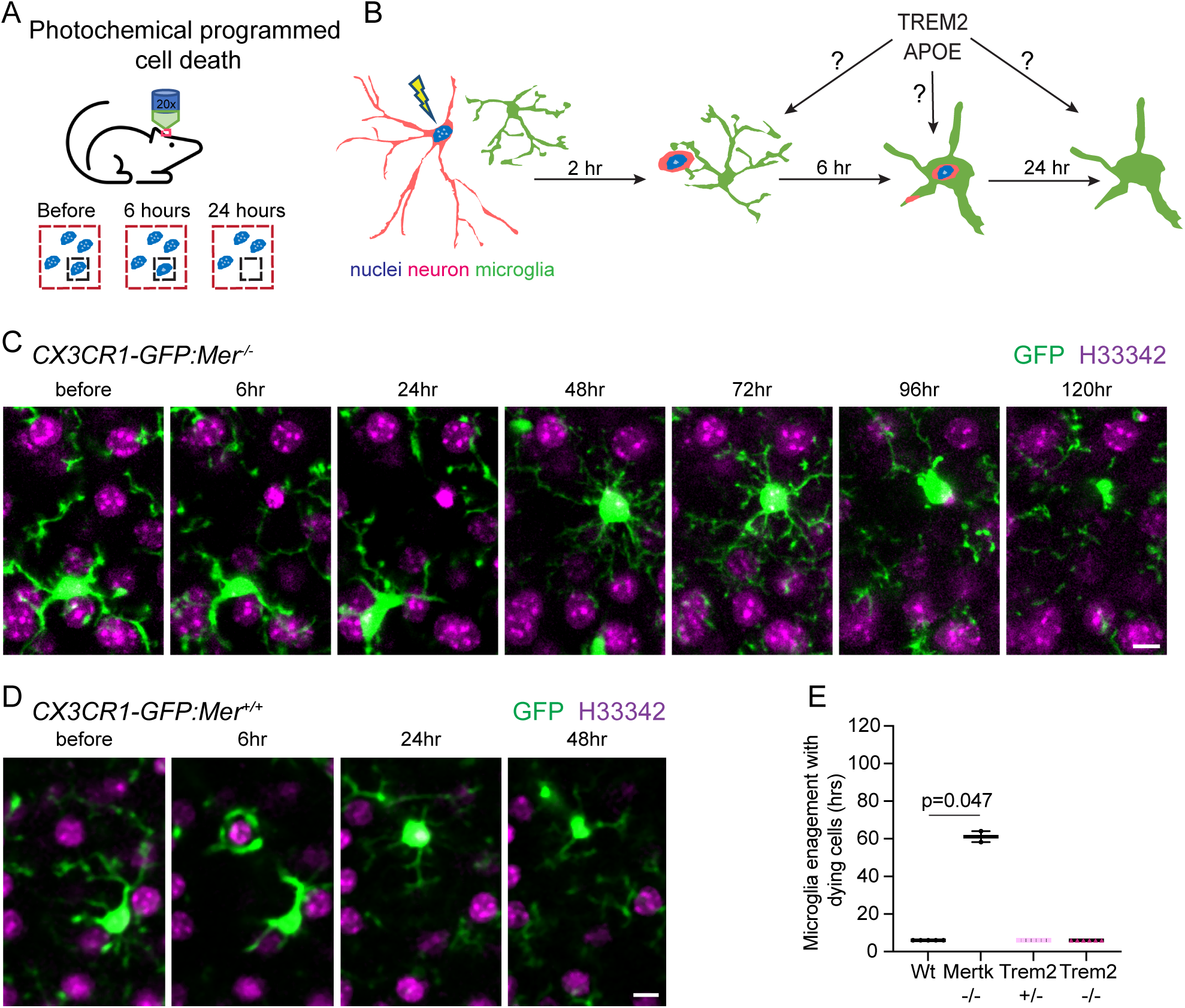
Strategy to investigate the molecular control of neuronal phagocytosis *in vivo*. (A) Schematic showing photochemical induction of cell death (2phatal) allowing visualization of dynamic nuclear changes in the live mouse brain and (B) examination of neuron and microglia interactions and testing of neurodegeneration related genes involved in corpse clearance. (C, D) Visualization of microglia (green) shows that there is a defective microglia engagement with dying cells (magenta) in *CX3CR1-GFP; Mer* ^*-/-*^ mouse leading to delayed corpse removal, as compared to rapid engagement and removal in *CX3CR1-GFP; Mer* ^*+/+*^ mice(D). Scale bar=10 µm. (E) Average time taken by microglia to engage with the neuronal corpse in wild type, Mertk^-/-^, Trem2 ^+/-^, and Trem2 ^-/-^. (n= 4 (Wild type); 2 (Mertk^-/-^); 4 (Trem2^+/-^); 6 (Trem2^-/-^) mice, and 10-15 cells per mouse).

### TREM2 or APOE deficiency do not impair phagocytosis of neuronal corpses *in vivo*

In AD, microglia exhibit a unique gene transcriptional profile that is believed to be regulated by TREM2 and APOE (Krasemann et al., 2017). In humans and mice, Trem2 and Apoe gene variants have been found to show significant deficiencies in microglia polarization toward amyloid plaques(Stephen et al., 2019; Wang et al., 2016; Yuan et al., 2016). We thus hypothesized that TREM2 or APOE deficiency could result in microglia polarization defects and consequent corpse clearance defects. To test this hypothesis, we applied our 2Phatal technique, inducing programmed cell death in the superficial cortical layers, and precisely analyzed the temporal course and efficiency of corpse removal in AD-like mice and mice with Trem2 or Apoe deletion. Interestingly, we found no differences either in the patterns of cell death and overall mortality (Fig 2A and Fig 2D (left panel)) or in the temporal course of corpse removal (Fig 2A and Fig 2D (right panel)) and microglia engagement with dying cells following 2Phatal induction (Fig 2B and Fig 1E). This finding suggests that in Trem2-deficient mice, microglia did not exert any detrimental effects on neurons nor impact corpse removal efficiency.

**Figure 2:**
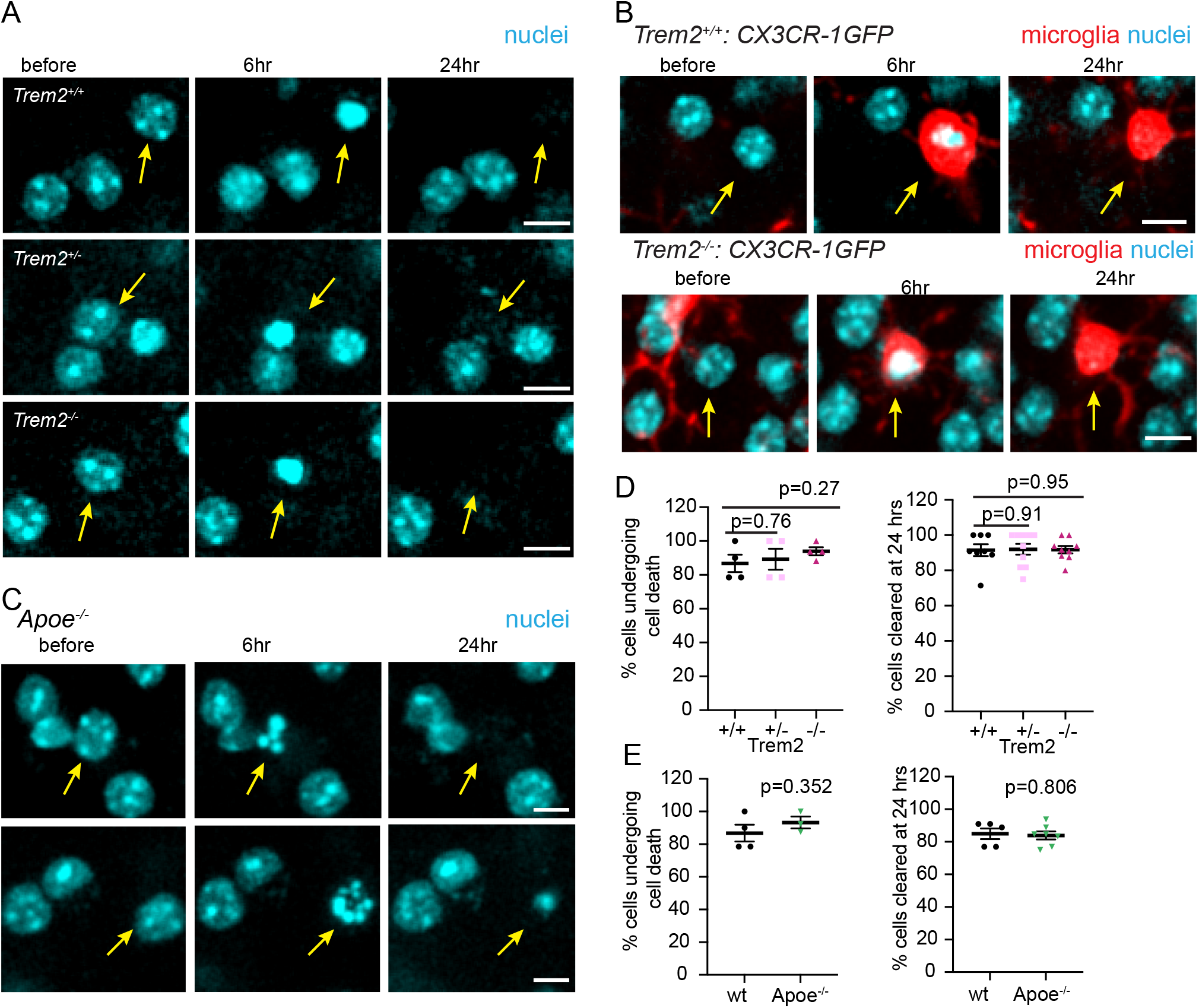
*Trem2* or *Apoe* deletion does not affect microglia engagement or phagocytosis of dying neurons. (A) *In vivo* time-lapse imaging of cell nucleus (cyan) undergoing condensation and removal during cell death (yellow arrow) in *Trem2* ^*+/+*^, *Trem2* ^*+/-*^, and *Trem2* ^*-/-*^ mice. (B) *In vivo* time lapse imaging of microglia (red) during corpse removal in *Trem2*^*+/+*^; *CX3CR1-GFP*^*+/-*^, and *Trem2*^*-/-*^; *CX3CR1-GFP*^*+/-*^ mice. In both *Trem2*^*+/+*^ and *Trem2*^*-/-*^ mice microglia migrated and engaged with the dying cells (yellow arrow) with equal efficiency. Scale bar= 10 µm. (C) *In vivo* time-lapse imaging of cell nucleus (cyan) undergoing condensation and removal during cell death (yellow arrow) in *Apoe*^*-/-*^ mice. Scale bar= 10 µm. (D) Quantification of cells undergoing cell death (left panel) and removal (right panel) (n= 4 to 10 mice in each group and 10-15 cells per mouse for each group) in *Trem2*^*+/+*^, *Trem2*^*+/-*^, and *Trem2*^*-/-*^. (E) Quantification of cells undergoing cell death (left panel) and removal (right panel) in *Apoe*^*-/-*^ mice (n= 5 to 7 mice in each group and 10-15 cells per animal).

Next, we proceeded to investigate whether Apoe played a role in regulating corpse removal *in vivo*. Like Trem2, we found that Apoe deficiency did not impact cell mortality rate or the efficiency of clearing dying neuron (Fig 2E & Fig 2F). To uncover any subtle effects of these genes in modulating corpse removal, we increased the phagocytic load by inducing the death of around 20 neurons simultaneously. Interestingly, we observed a marked reduction in overall corpse removal efficiency under these conditions (Fig 3A & Fig 3B). However, in Trem2 or Apoe-deficient mice, we did not observe any additional delay in corpse removal when we performed the same experiment (Fig 3C & Fig 3D). In conclusion, our results suggest that under normal conditions TREM2 and APOE do not play a significant role in regulating microglia behavior during corpse clearance.

**Figure 3:**
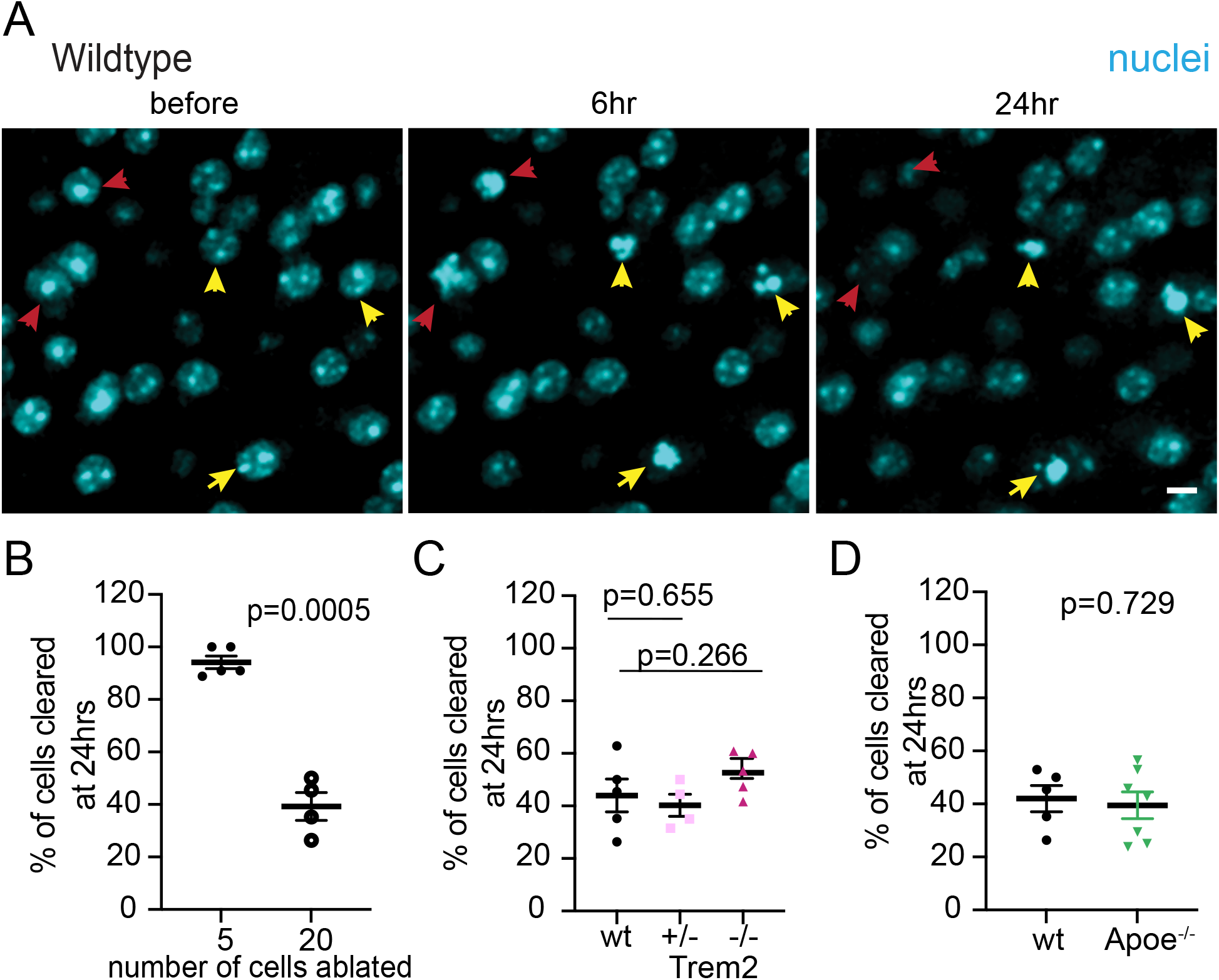
Increasing cell death load reduces phagocytic efficiency in a TREM2 and APOE-independent manner. (A) *In vivo* time lapse images showing nuclear condensation and clearance in wildtype mice when cell death was induced in more than 20 cells in a region of interest. Cells that were removed by 24 hrs are shown with red arrow heads and those persistent are shown in yellow arrows. Scale bar= 10 µm (B) Percentage of cell cleared at 24 hrs in wild type mice with low (n=5 mice and 10-15 cells per mouse) and high cell death load (n=4 mice and 50-60 cells per mouse). (C,D) Percentage of cells cleared at 24 hours following large cell death load in wild type, *Trem2*^*+/-*^, *Trem2* ^*-/-*^, and *Apoe* ^*-/-*^ mice (n=4 to 7 mice per group and 50-60 cells per mouse).

### Corpse removal in Alzheimer-like mice is not dependent on Trem2 function

In AD, Trem2 has been identified as a critical factor in the chronic polarization of microglia towards amyloid plaques (Yuan et al., 2016). Previous studies have shown that Trem2 also regulates the metabolic fitness (Ulland et al., 2017) and transition to a late-stage DAM profile (DAM2) (Keren-Shaul et al., 2017) of microglia. Given its known role in these processes, we hypothesized that we could observe a change in corpse clearance efficiency in Trem2-deficient mice when dying cells were in close proximity to extracellular amyloid aggregates. Surprisingly, our results showed that Trem2 deficiency did not have any significant effect on the phagocytic clearance of dying cells, regardless of their proximity to the plaque edges (Fig 4A-C). Even when we further increased the phagocytic load by inducing the death of a larger number of cells, resulting in a marked reduction in corpse clearance efficiency (Fig 3B), Trem2 haploinsufficiency did not further affect the efficiency of corpse removal (Fig 4D). These findings suggest that despite the prominent phenotypic changes of microglia around plaques and their dependence on Trem2 for polarization, they still retain normal phagocytic capabilities in the absence of Trem2.

**Figure 4:**
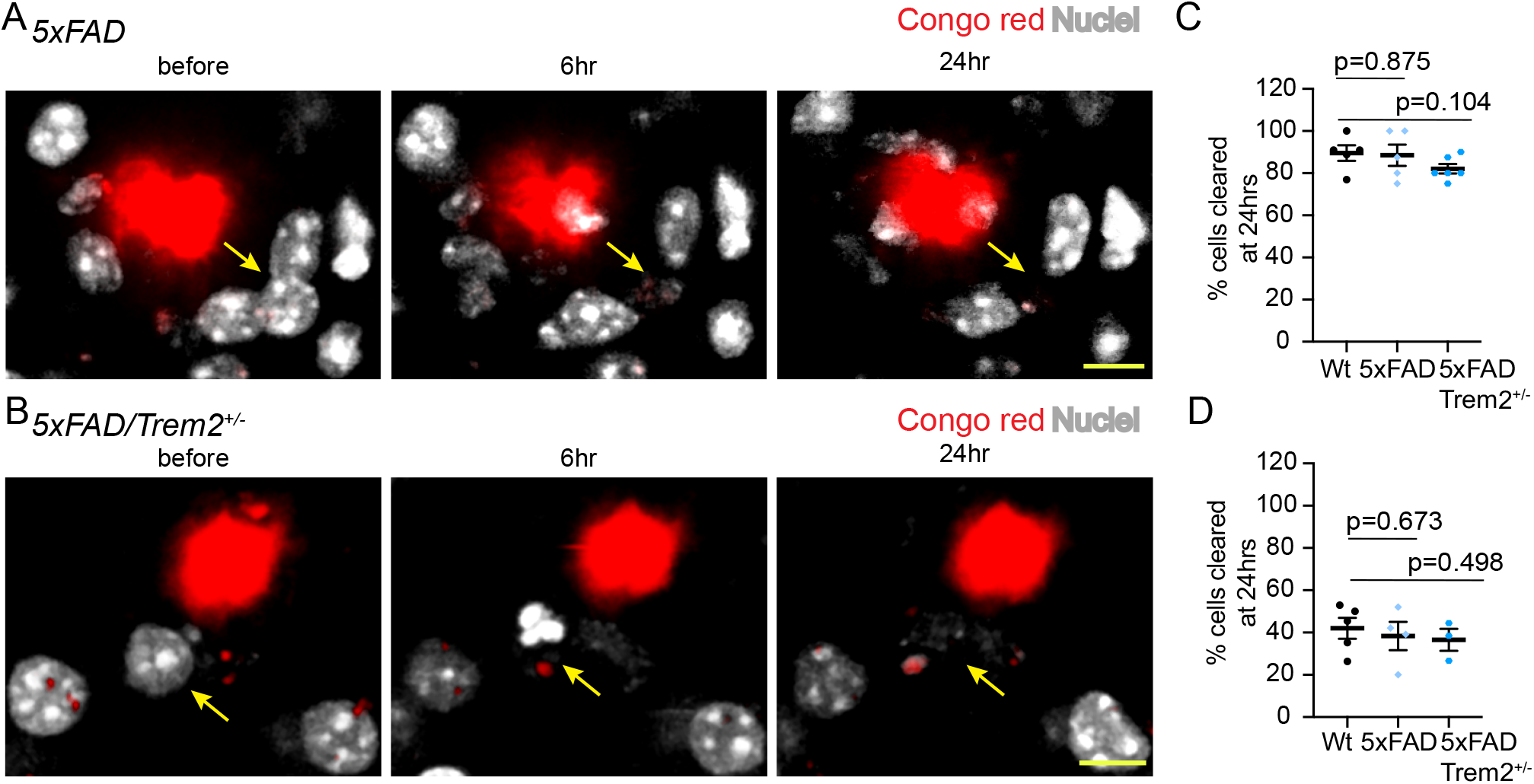
Phagocytic corpse clearance by microglia is not affected by the presence of amyloid plaques. (A, B) *In vivo* time lapse imaging of dying cells (yellow arrow) and plaques (red) in *5xFAD* and *5xFAD/Trem2*^*+/-*^ mice. Scale bar= 10 µm (C) Percentage of cells cleared at 24 hours following low (n= 3 to 6 mice per group and 10-15 cells per mouse) and high cell death load (D) (n= 3 to 6 mice per group and 50-60 cells per mouse) in Wild type, *5xFAD*, and *5xFAD/Trem2*^*+/-*^.

### Plaque-associated microglia become migratory during corpse removal in the absence of Trem2

Previous studies have reported that in Trem2-deficient mice, microglia processes are unable to tightly wrap around the amyloid plaque surface (Yuan et al., 2016). To further investigate the dynamic behavior of microglia around plaques following programmed cell death induction, we visualized microglia using triple mutant mice (5xFAD/Trem2+/-/CX3CR1-GFP+/-). Our findings show that in Trem2 wild type mice, microglia with robust process polarization towards the plaque surface were capable of phagocytosing adjacent dying cells by projecting engulfing processes towards them (Fig 5A). Interestingly, their strong polarization towards plaques was maintained throughout the entire phagocytic process. In contrast, in Trem2 haploinsufficiency, microglia displayed processes that projected towards plaques, but were not closely associated with them. However, they were still able to phagocytose dying cells in their vicinity. They did so by rapidly moving their cell bodies towards the dying cells, during which their attachment to plaques was further lost (Fig 5B). These findings suggest that while Trem2 has no impact on the engagement with dying cells (Fig 5C) or efficiency of corpse phagocytosis (Fig 4C), it has a dramatic effect on the anchoring of microglia processes towards plaques and the adoption of a more migratory phenotype during corpse removal, similar to what is observed with microglia in wild type mice.

**Figure 5:**
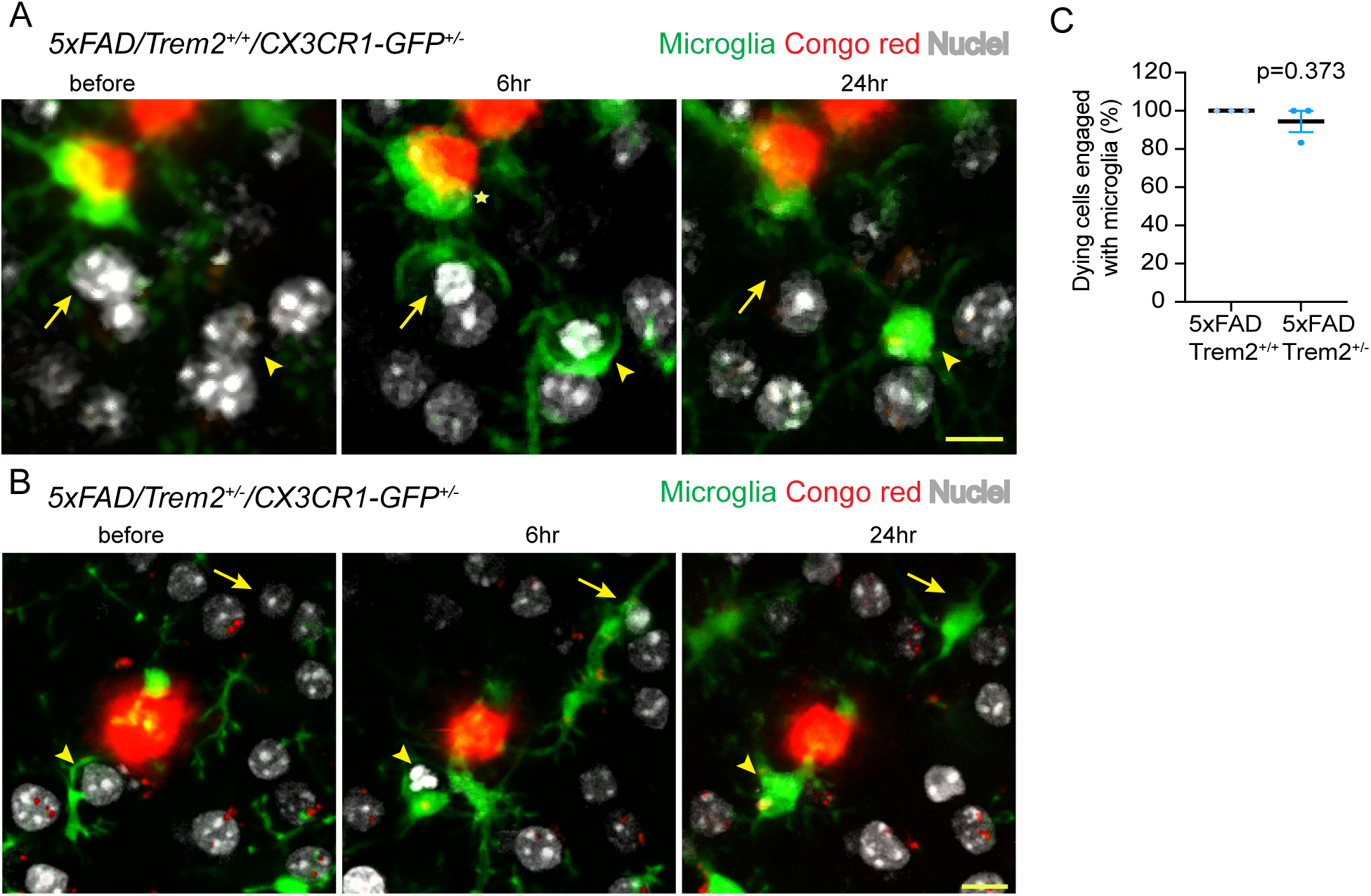
Trem2 deletion leads microglia to prioritize corpse engulfment over plaque encapsulation. (A, B) *In vivo* time lapse imaging of microglia engagement with the dying cells in *5xFAD/Trem2*^*+/+*^*/CX3CR1-GFP*^*+/-*^ and *5xFAD/Trem2*^*+/-*^*/CX3CR1-GFP*^*+/-*^. In *5xFAD/Trem2*^*+/+*^ mice, a microglia (green, asterisk) that is engaged with the plaque (red) extended its processes towards the nearby dying cells (yellow arrow) while maintaining its amyloid-facing processes polarized towards the plaque surface. However, in *5xFAD/Trem2*^*+/-*^ mice (B) microglia were less polarized towards the amyloid plaques (red), and readily migrated their cell bodies to engulf neuronal corpses. Scale bar= 10 µm. (C) Proportion of dying cells engaged with microglia in *5xFAD/Trem2*^*+/+*^*/CX3CR1-GFP*^*+/-*^ and *5xFAD/Trem2*^*+/-*^*/CX3CR1-GFP*^*+/-*^ mice (n=3 mice per group and 3 to 5 cells per mouse).

## DISCUSSION

In this study, we used innovative intravital techniques to examine cell death and corpse removal in order to measure the effect of neurodegeneration risk genes on phagocytosis efficiency in the brain. Our focus was on two genes: *Trem2* and *Apoe*. Previous research has linked these genes to microglia polarization, phagocytosis of amyloid deposits, and the adoption of a DAM phenotype (Keren-Shaul et al., 2017; Kleinberger et al., 2014; Krasemann et al., 2017; Yuan et al., 2016). Additionally, Trem2 has been found to play a role in injury responses and the clearance of apoptotic cells by microglia through the sensing of aminophospholipids (Hsieh et al., 2009; Shirotani et al., 2019; Takahashi et al., 2005). Apoe, which is highly expressed in microglia (Rodriguez et al., 2014) was recently suggested to act as a potential ligand for Trem2 (Atagi et al., 2015), and also to regulate the switch to a DAM phenotype, ultimately affecting microglia polarization and amyloid phagocytosis (Krasemann et al., 2017).

Previous research has generated conflicting evidence about the role of Trem2 in phagocytosis. Studies conducted in cell cultures assaying phagocytosis of apoptotic cells and beta-amyloid aggregates have produced conflicting results, with some indicating no deficiencies and others reporting phagocytic defects resulting from Trem2 deletion (Cignarella et al., 2020; Poliani et al., 2015; Wang et al., 2015; Wang et al., 2023). *In vivo* studies in mice with Trem2 deficiency have only showed subtle differences in myelin phagocytosis and remyelination following extensive demyelination (Hsieh et al., 2009; Takahashi et al., 2007; Takahashi et al., 2005; Wang et al., 2015). However, our *in vivo* studies, conducted quantitatively in the live brain, at single cell resolution, provided strong evidence that neither TREM2 nor APOE plays a significant role in detecting, engulfing, or digesting neuronal corpses.

Although Trem2 and Apoe have been implicated in controlling the evolution of a disease-associated microglia (DAM) state in AD-like mice (Keren-Shaul et al., 2017; Krasemann et al., 2017), we did not observe any defects in corpse removal with deletion of these genes. Our findings suggest that Trem2 and Apoe may mediate microglial responses in chronically polarized states, such as towards amyloid plaques, rather than in acute responses such as those observed after cell death. Additionally, our study revealed that once microglia are chronically polarized towards plaques, their processes and cell bodies do not disengage or migrate towards neuronal corpses. However, in the absence of TREM2 signaling, we observed that plaque-facing microglia can rapidly disengage and migrate towards dying cells. This indicates that TREM2 plays a critical role in chronic polarization of microglia, but not in their ability to migrate during phagocytosis. This observation suggests that chronic encapsulation of foreign material, parasites, or other infectious agents could have been evolutionarily prioritized over homeostatic functions such as corpse removal.

There are several factors that may account for the discrepancies between our findings and previous studies. First, it is possible that differences in experimental methods could contribute to conflicting results. While previous investigations primarily utilized microglia-like cells exposed to high levels of dying cells in vitro, our in vivo model more accurately reflects programmed cell death and glial corpse clearance in the brain. Second, our methodology allows for precise visualization and quantitative analysis of individual dying neurons and subsequent microglial engagement and clearance, which provides a more detailed understanding of the process compared to other complex in vivo injury models. Third, our techniques are highly sensitive, as demonstrated by our ability to identify clear phenotypes in other gene deletions such as MerTK, which is well-known to be involved in phagocytosis signaling. Moreover, we have observed a phenotype of slowed microglial engagement and phagocytosis in situations where a larger number of neurons are killed (Fig 3A & Fig 3B) or in aging mice (Damisah et al., 2020), supporting the validity of our methods.

One limitation of our study is that we only investigated neuronal death but not death of other cell types, and there could be cell-type specific differences in the dynamics of cell death, glial engulfment (Damisah et al., 2020; Hill et al., 2017) and the roles of modulating genes. Indeed, several studies have reported potential roles of Trem2 in myelin phagocytosis(Cignarella et al., 2020). It is thus possible that unlike in neuronal death, phagocytosis during myelin shedding in aging and pathology may be modulated by different signaling pathways, including TREM2 and APOE. In the future, it would be interesting to examine this using our methodologies combined with intravital imaging of myelin and oligodendrocytes (Schain et al., 2014).

An additional limitation is that our study used relatively young mice and in theory a delayed phagocytosis phenotype difference between Trem2 and Apoe knockout mice and controls could be uncovered with aging. Finally, while our 2Phatal method induces a form of programmed cell death, the precise mechanisms are still unclear, and they may differ from naturally occurring apoptosis or necroptosis. We can thus not rule out that TREM2 and APOE could have more prominent roles in modulating glial engulfment in such forms of cell death.

Overall, our studies shed light on the cellular and molecular mechanisms governing the removal of dying cells by microglia in both normal and disease states. The novel methodologies we developed could prove useful in future studies investigating other genes and molecules involved in phagocytosis, and for testing potential therapeutic interventions. Ultimately, a deeper understanding of the complex interplay between glial cells and dying neurons could have important implications for the development of treatments for neurodegenerative diseases.

## METHODS

### Transgenic mouse strains

In this study, we used several genetically modified mouse models, including Trem2 KO (JAX stock#027197), Apoe KO (JAX stock #002052), 5xFAD (JAX stock#34840), and CX3CR1-GFP (JAX stock#005582). To visualize microglia engagement, Trem2 KO mice were crossbred with CX3CR1-GFP mice. To study the effect on phagocytosis of dying cells by polarized microglia in AD, 5xFAD mouse were crossbred with CX3CR1-GFP mouse. Finally, a triple transgenic mouse was created by crossbreeding 5xFAD with Trem2-/-;CX3CR1-GFP to visualize phagocytosis by microglia in AD with Trem2 haploinsufficiency. In all experiments mice of either sex was used. All animal procedures were approved by the Institutional Animal Care and Use committee at Yale University.

### Cranial window surgery and in vivo imaging

The cranial window implant and dye application procedure was conducted following established protocols (Damisah et al., 2020; Hill et al., 2017). Briefly, mice were anesthetized with intraperitoneal injections of Ketamine and xylazine solutions (100 mg per kg and 10 mg per kg, respectively). The scalp was shaved, and a small piece of skin was removed to expose the skull. A 3mm diameter circle was drilled, and fine forceps were used to remove the dura without damaging the brain surface. Hoechst 33342 (H3570 Thermo Fischer Scientific) and/or Congo red (Sigma-Aldrich C6277) were diluted in PBS and applied topically for 5-10 minutes. A glass coverslip was gently pressed and glued to the skull. Post-surgery, mice were placed on a heating pad for recovery.

The in vivo imaging was performed using a previously published technique (Damisah et al., 2020). In brief, the mice were anesthetized and positioned under a two-photon microscope that was equipped with a mode-locked MaiTai tunable laser from Spectra Physics. The fluorophores were excited at specific wavelengths: Hoechst 33342 at 775nm, GFP and Congo red at 900nm. For each region of interest (ROI), a landmark image capturing the large blood vessels was recorded and used for longitudinal imaging.

### Photochemical programmed cell death

Induction of programmed cell death was performed as previously described (Damisah et al., 2020; Hill et al., 2017) using two-photon laser excitation. Briefly, the laser was fixed at 775 nm and a 20×20 pixel region of interest centered at the targeted cell nuclei labeled with Hoechst 33342 was selected. Photobleaching was conducted using a dwell time of 100 microseconds and a power of 20-40 mW for 10 seconds. Images were captured before and after photobleaching, as well as at 6 and 24 hours post-bleaching to monitor changes in cell nuclei, microglial engagement, and phagocytosis. The short duration of photochemical bleaching and low laser power ensured that there was no thermal cellular injury.

### Image processing and quantification

The displayed images (Figure 1-5) depict the maximal optical projection of three original XYZ slices centered at the middle plane of the selected nuclei. The images were processed using the Fiji/Image J software, and the same procedure was applied to all time points. The Despeckle process tool was used to remove noise, and this was applied to all time points. To account for the non-specific binding of Hoechst dye to the plaques, we subtracted the Hoechst channel from the Congo red channel for each time-point (Fig4).

To quantify the percentage of cells undergoing cell death in Trem2 and Apoe knockout mice, the condensation of neuronal cell nuclei post-photo bleaching at the 6-hour time-point was manually counted. Each data point represents the percentage of the total number of cell nuclei condensed by the total number of cells photo-bleached per animal (Fig2b, Fig2e). Similarly, the absence of neuronal nuclei at 24 hours was manually counted to record for clearance by microglia in each genotype (Fig 2b, 2e, 3b, 3c, 3d, 4b, and 5b). Microglia engagement was quantified by recording the microglia phagocytic cup around the condensed nuclei at 6 hours (Fig 1E and Fig 5C). The graph representing the time required for microglia to engage with the dying cell in MerTk knock - out mice is a reanalysis of previously collected data (Damisah et al., 2020).

### Statistics

Statistical analysis was conducted using Prism software. Unpaired Student’s t-test (non-parametric) with Welch’s correction was utilized to determine the statistical significance of the data represented in Fig 1-5. Individual data points and standard error mean is represented in the graphs. P value for each comparison is mentioned in the graph itself. Normal distribution and equal variance were assumed for all data in each statistical test. No data were excluded from the analysis, randomization was not used to assign experimental subjects, and experimenter blinding was not required. The sample size for the experiments was not predetermined using any statistical methods. The figure legends indicate the number of animals used in each experiment.

## Data Availability

All data necessary to evaluate the conclusions in the paper are included in the paper. Any additional data that may be required can be obtained from the authors upon reasonable request.

## Author Contributions

E.C.D, R.A.H. and J.G. conceived of the study. E.C.D., A.R, R.A.H. designed all experiments. E.C.D., A.R, R.A.H., and L.H. performed all experiments E.C.D. A.R and J.G. wrote the manuscript. J.G. directed the study.

## References

Atagi, Y., C.C. Liu, M.M. Painter, X.F. Chen, C. Verbeeck, H. Zheng, X. Li, R. Rademakers, S.S. Kang, H. Xu, S. Younkin, P. Das, J.D. Fryer, and G. Bu. 2015. Apolipoprotein E Is a Ligand for Triggering Receptor Expressed on Myeloid Cells 2 (TREM2). J Biol Chem 290:26043–26050.

Cignarella, F., F. Filipello, B. Bollman, C. Cantoni, A. Locca, R. Mikesell, M. Manis, A. Ibrahim, L. Deng, B.A. Benitez, C. Cruchaga, D. Licastro, K. Mihindukulasuriya, O. Harari, M. Buckland, D.M. Holtzman, A. Rosenthal, T. Schwabe, I. Tassi, and L. Piccio. 2020. TREM2 activation on microglia promotes myelin debris clearance and remyelination in a model of multiple sclerosis. Acta Neuropathol 140:513–534.

Damisah, E.C., R.A. Hill, A. Rai, F. Chen, C.V. Rothlin, S. Ghosh, and J. Grutzendler. 2020. Astrocytes and microglia play orchestrated roles and respect phagocytic territories during neuronal corpse removal in vivo. Sci Adv 6:eaba3239.

Gal, A., Y. Li, D.A. Thompson, J. Weir, U. Orth, S.G. Jacobson, E. Apfelstedt-Sylla, and D. Vollrath. 2000. Mutations in MERTK, the human orthologue of the RCS rat retinal dystrophy gene, cause retinitis pigmentosa. Nat Genet 26:270–271.

Hill, R.A., E.C. Damisah, F. Chen, A.C. Kwan, and J. Grutzendler. 2017. Targeted two-photon chemical apoptotic ablation of defined cell types in vivo. Nat Commun 8:15837.

Hsieh, C.L., M. Koike, S.C. Spusta, E.C. Niemi, M. Yenari, M.C. Nakamura, and W.E. Seaman. 2009. A role for TREM2 ligands in the phagocytosis of apoptotic neuronal cells by microglia. J Neurochem 109:1144–1156.

Keren-Shaul, H., A. Spinrad, A. Weiner, O. Matcovitch-Natan, R. Dvir-Szternfeld, T.K. Ulland, E. David, K. Baruch, D. Lara-Astaiso, B. Toth, S. Itzkovitz, M. Colonna, M. Schwartz, and I. Amit. 2017. A Unique Microglia Type Associated with Restricting Development of Alzheimer’s Disease. Cell 169:1276–1290 e1217.

Kleinberger, G., Y. Yamanishi, M. Suarez-Calvet, E. Czirr, E. Lohmann, E. Cuyvers, H. Struyfs, N. Pettkus, A. Wenninger-Weinzierl, F. Mazaheri, S. Tahirovic, A. Lleo, D. Alcolea, J. Fortea, M. Willem, S. Lammich, J.L. Molinuevo, R. Sanchez-Valle, A. Antonell, A. Ramirez, M.T. Heneka, K. Sleegers, J. van der Zee, J.J. Martin, S. Engelborghs, A. Demirtas-Tatlidede, H. Zetterberg, C. Van Broeckhoven, H. Gurvit, T. Wyss-Coray, J. Hardy, M. Colonna, and C. Haass. 2014. TREM2 mutations implicated in neurodegeneration impair cell surface transport and phagocytosis. Sci Transl Med 6:243ra286.

Krasemann, S., C. Madore, R. Cialic, C. Baufeld, N. Calcagno, R. El Fatimy, L. Beckers, E. O’Loughlin, Y. Xu, Z. Fanek, D.J. Greco, S.T. Smith, G. Tweet, Z. Humulock, T. Zrzavy, P. Conde-Sanroman, M. Gacias, Z. Weng, H. Chen, E. Tjon, F. Mazaheri, K. Hartmann, A. Madi, J.D. Ulrich, M. Glatzel, A. Worthmann, J. Heeren, B. Budnik, C. Lemere, T. Ikezu, F.L. Heppner, V. Litvak, D.M. Holtzman, H. Lassmann, H.L. Weiner, J. Ochando, C. Haass, and O. Butovsky. 2017. The TREM2-APOE Pathway Drives the Transcriptional Phenotype of Dysfunctional Microglia in Neurodegenerative Diseases. Immunity 47:566–581 e569.

Moujalled, D., A. Strasser, and J.R. Liddell. 2021. Molecular mechanisms of cell death in neurological diseases. Cell Death Differ 28:2029–2044.

Poliani, P.L., Y. Wang, E. Fontana, M.L. Robinette, Y. Yamanishi, S. Gilfillan, and M. Colonna. 2015. TREM2 sustains microglial expansion during aging and response to demyelination. J Clin Invest 125:2161–2170.

Rademakers, R., M. Baker, A.M. Nicholson, N.J. Rutherford, N. Finch, A. Soto-Ortolaza, J. Lash, C. Wider, A. Wojtas, M. DeJesus-Hernandez, J. Adamson, N. Kouri, C. Sundal, E.A. Shuster, J. Aasly, J. MacKenzie, S. Roeber, H.A. Kretzschmar, B.F. Boeve, D.S. Knopman, R.C. Petersen, N.J. Cairns, B. Ghetti, S. Spina, J. Garbern, A.C. Tselis, R. Uitti, P. Das, J.A. Van Gerpen, J.F. Meschia, S. Levy, D.F. Broderick, N. Graff-Radford, O.A. Ross, B.B. Miller, R.H. Swerdlow, D.W. Dickson, and Z.K. Wszolek. 2011. Mutations in the colony stimulating factor 1 receptor (CSF1R) gene cause hereditary diffuse leukoencephalopathy with spheroids. Nat Genet 44:200–205.

Rodriguez, G.A., L.M. Tai, M.J. LaDu, and G.W. Rebeck. 2014. Human APOE4 increases microglia reactivity at Abeta plaques in a mouse model of Abeta deposition. J Neuroinflammation 11:111.

Schain, A.J., R.A. Hill, and J. Grutzendler. 2014. Label-free in vivo imaging of myelinated axons in health and disease with spectral confocal reflectance microscopy. Nat Med 20:443–449.

Shirotani, K., Y. Hori, R. Yoshizaki, E. Higuchi, M. Colonna, T. Saito, S. Hashimoto, T. Saito, T.C. Saido, and N. Iwata. 2019. Aminophospholipids are signal-transducing TREM2 ligands on apoptotic cells. Sci Rep 9:7508.

Stephen, T.L., M. Cacciottolo, D. Balu, T.E. Morgan, M.J. LaDu, C.E. Finch, and C.J. Pike. 2019. APOE genotype and sex affect microglial interactions with plaques in Alzheimer’s disease mice. Acta Neuropathol Commun 7:82.

Takahashi, K., M. Prinz, M. Stagi, O. Chechneva, and H. Neumann. 2007. TREM2-transduced myeloid precursors mediate nervous tissue debris clearance and facilitate recovery in an animal model of multiple sclerosis. PLoS Med 4:e124.

Takahashi, K., C.D. Rochford, and H. Neumann. 2005. Clearance of apoptotic neurons without inflammation by microglial triggering receptor expressed on myeloid cells-2. J Exp Med 201:647–657.

Ulland, T.K., W.M. Song, S.C. Huang, J.D. Ulrich, A. Sergushichev, W.L. Beatty, A.A. Loboda, Y. Zhou, N.J. Cairns, A. Kambal, E. Loginicheva, S. Gilfillan, M. Cella, H.W. Virgin, E.R. Unanue, Y. Wang, M.N. Artyomov, D.M. Holtzman, and M. Colonna. 2017. TREM2 Maintains Microglial Metabolic Fitness in Alzheimer’s Disease. Cell 170:649–663 e613.

Vila, M., and S. Przedborski. 2003. Targeting programmed cell death in neurodegenerative diseases. Nat Rev Neurosci 4:365–375.

Wang, Y., M. Cella, K. Mallinson, J.D. Ulrich, K.L. Young, M.L. Robinette, S. Gilfillan, G.M. Krishnan, S. Sudhakar, B.H. Zinselmeyer, D.M. Holtzman, J.R. Cirrito, and M. Colonna. 2015. TREM2 lipid sensing sustains the microglial response in an Alzheimer’s disease model. Cell 160:1061–1071.

Wang, Y., R.V. Kyauk, Y.A. Shen, L. Xie, M. Reichelt, H. Lin, Z. Jiang, H. Ngu, K. Shen, J.J. Greene, M. Sheng, and T.J. Yuen. 2023. TREM2-dependent microglial function is essential for remyelination and subsequent neuroprotection. Glia

Wang, Y., T.K. Ulland, J.D. Ulrich, W. Song, J.A. Tzaferis, J.T. Hole, P. Yuan, T.E. Mahan, Y. Shi, S. Gilfillan, M. Cella, J. Grutzendler, R.B. DeMattos, J.R. Cirrito, D.M. Holtzman, and M. Colonna. 2016. TREM2-mediated early microglial response limits diffusion and toxicity of amyloid plaques. J Exp Med 213:667–675.

Yuan, P., C. Condello, C.D. Keene, Y. Wang, T.D. Bird, S.M. Paul, W. Luo, M. Colonna, D. Baddeley, and J. Grutzendler. 2016. TREM2 Haplodeficiency in Mice and Humans Impairs the Microglia Barrier Function Leading to Decreased Amyloid Compaction and Severe Axonal Dystrophy. Neuron 90:724–739.

